# Animals exhibit personality traits in their movement: a case study on location trajectories of primates

**DOI:** 10.1101/775486

**Authors:** Takashi Morita, Aru Toyoda, Seitaro Aisu, Akihisa Kaneko, Naoko Suda-Hashimoto, Ikki Matsuda, Hiroki Koda

## Abstract

Researching individual recognition (IR) is essential to understand the life history and adaptive behavior of social animals. Investigation of personality traits may also provide insights into how social animals distinguish between different individuals. This study investigates IR behavior in Japanese macaques (*Macaca fuscata*), focusing on one specific trait, which is movement. Using a recently developed tracking system based on Bluetooth^®^ Low Energy beacons, we collected three-dimensional (3D) location data from five Japanese macaques living in a group cage. A non-parametric, neural network-based analysis of the data revealed the existence of personality traits in extremely limited aspects of the movement data (2-min trajectory of 3D location). Our results support the validity of multimodal approaches in studying IR, beyond the typical single-frame face recognition method, both for researchers and animal agents.

## 1 Introduction

Individual recognition (IR) is an important component in the behavioral ecology of social animals, influencing an individuals’ life history characteristics and broader species and inclusive fitness [1]. Researchers have investigated a wide variety of personality, genetic, and physical traits involved in IR—including facial features, body coloration, vocalizations, and chemical signals—and recent achievements in machine learning have allowed more large-scale analysis [2, 3]. Personality traits have also been discussed in the context of cognitive modeling: how do social animals themselves identify one another in their group [4, 5]?

This study investigates movement as a personality trait *movement* because previous research has shown that human body movement is significantly influenced by personality, even in simple movements like point-light walking, which may not involve personality traits like face/body shapes [6, 7]. We specifically study another limited aspect of movement, namely *location trajectory*. Given the importance of this personality trait in humans-to the point where this information is a central point of privacy (e.g., GPS tracking by smartphones), we also consider it to be an important personality trait in non-human primates. We present a case study of Japanese macaques (*Macaca fuscata*) in captivity herein.

Japanese macaques have often been employed in comparative studies [8]—for instance, their social hierarchy and philopatry show various similarities to/differences from human society—and have been widely studied for better understanding of the evolution of social systems. IR is considered as a crucial behavioral trait in such social species, where individuals need to adapt their behavior in relation to different group members. This study contributes to the present body of research, introducing a novel data collection and analysis methodology and providing a baseline result for future studies on human and other social species, particularly those lacking traditional IR mechanisms such as face recognition.

## 2 Materials & Methods

### 2.1 Ethical Statement

All procedures were reviewed and approved by the Animal Welfare and Care Committee of the Primate Research Institute, Kyoto University (KUPRI; Permission # 2018-203) and complied with the institutional guidelines [9].

### 2.2 Subjects and Facilities

We conducted observations of five adult Japanese macaques (two males and three females) in an outdoor cage (4 × 5 × 2.5 m) installed in the KUPRI, Japan. The subjects were allowed to freely move in the cage and were exposed to outside temperatures—with natural fluctuations—throughout the study. Following the standard nutritional requirements for non-human primates, the animals were fed primate pellets with supplemental vegetables (sweet potato) once a day.

### 2.3 Data Collection

We recorded three-dimensional (3D) location trajectories of five individuals using a recently developed real-time tracking system based on Bluetooth^®^ Low Energy (BLE) beacons. Each individual macaque carried two BLE beacons (Quuppa Intelligent Locating System™) attached to a custom-made collar (see Figure S1.1). The subjects were collared on the first day of the study and the collars were removed on the last day. The collaring process was as follows: first, the subjects were separated from their original group and held in a designated area; they were then immobilized with ketamine (10 mg/kg intramuscular injection) and collared. The collar was made to fit each individual, neither too loosely (to reduce shaking of beacons) nor too tightly (to minimize subject discomfort and prevent injury). According to the recommended institutional guidelines (i.e. 5% of the subjects’ body weight) [9], the collars weighed approximately 40 gm each. Once collared, the subjects were awoken with an injection of approximately 1,200 *µ*g/kg atipamezole.

We installed six receivers for the beacon signals: four above and two on the sides of the cage. The system sampling frequency was set at 9 Hz, although the actual rate was unstable because of uncontrollable signal interference and reflection perturbation.

We analyzed the data collected between December 6 and 19 2018. Observations started an hour after collaring and ended an hour before removing the collars. Individual observations were split into 2-min intervals with a 30-s pause between the onsets, resulting in 175,494 time series. These data defined the location trajectories from which we assessed IR behavior of the individual macaques.

### 2.4 Data Analysis

As stated above, we intended to investigate to what extent individual macaques are identifiable from the trajectory of their location (i.e. movement). We measured the probability ℙ(*I* = *i* | **s**) of each individual’s ID number *i* ∈ {1, …, 5} conditioned on the time series of 3D coordinates, 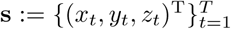, representing the trajectory of an individual’s location (not necessarily of *i*) in a 2-min time interval (including *T* data points). There is a lack of research on the links between movement and identity of individual primates (or other animals), and very little information on the most appropriate analysis for the conditional probability mass function ℙ(· | ·). Accordingly, we trained a neural network to obtain an optimal estimation of ℙ(· | ·) using 90% of the data (see S2 for details on the neural network and S3 for the data size per individual).

The trained network was used to compute the accuracy of the predictions of the remaining 10% of the data. We also measured the accuracy limiting the ground truth individual to each subject: that is, we reported the *recall* scores with each subject as the target. High recall scores indicate few false negatives and, thus, a great amount of personality information included in the location trajectories of the target individual. S5 reports all scores (including recall scores) in the standard machine learning evaluation, including precision and F1 scores as supporting information (see S4 for the mathematical definition of the scores). All scores had a chance-level expectation of 5^−1^ = 0.2, and we measured their statistical significance by comparing 100,000 bootstrapped scores against this chance level [10].

## 3 Results

Table 1 reports the model predictions from the test data. The model scored significantly greater overall accuracy than the chance level (*p* < 0.001), indicating that the macaques exhibited personality in their location trajectories. The model also predicted each individual with a significantly greater recall score than the chance level (*p* < 0.001; the precision and F1 scores, reported in S5 were also significantly larger), indicating that every subject—not just some of them—exhibited characteristic location trajectories.

**Table 1:** Scores of the model predictions with the statistical significance against the chance level (= 0.2), estimated from 100,000 bootstrapped samples.

| Target Individual | Score Type | Value | 95%CI | vs. Chance |
| --- | --- | --- | --- | --- |
| Overall | Accuracy | 0.372308 | [0.365185, 0.379430] | $p < 0.001$ |
| Female 1 | Recall | 0.247166 | [0.232932, 0.261500] | $p < 0.001$ |
| Female 2 | | 0.318512 | [0.303320, 0.333883] | $p < 0.001$ |
| Female 3 | | 0.301684 | [0.286374, 0.316973] | $p < 0.001$ |
| Male 1 | | 0.577461 | [0.560996, 0.593891] | $p < 0.001$ |
| Male 2 | | 0.417501 | [0.401249, 0.433796] | $p < 0.001$ |

The recall scores also showed variability between subjects: individuals were ranked as Male 1 > Male 2 > Female 2 > Female 3 > Female 1, and the 95% confidence intervals of the scores (estimated from the 100,000 bootstrapped samples) did not overlap, except between Female 2 and Female 3. As we can see, the males yielded greater recall scores than the females.

## 4 Discussion

To our knowledge, the significant accuracy and recall scores reported here provide the first evidence of the influence of personality traits in the location trajectory of non-human primates. Although the scores were not significant in terms of biometrics (e.g. > 90% accuracy with face recognition [2]), movement-based identification would still be useful to study behavioral ecology, particularly when state-of-the-art identification methods are not applicable. This method also enables comparative studies for a broader range of species with different physical characteristics. Additionally, previous studies on machine learning have shown that multimodal data often leads to more successful outcomes [11, 12, 13]. Personality traits in movement may contribute to better IR, particularly in combination with other techniques such as face recognition.

Male macaques exhibited greater recall scores—indicating a greater degree of personality—than females. Given that the sample size (*n* = 5) of this study was small, we caution that the results obtained here may simply be a result of individual variation. However, the reported gender-based difference in scores could be due to differences in the social traits: for instance, female Japanese macaques are known to exhibit greater spatial cohesion than males [14]. Furthermore, macaques tend to confine their grooming to kin, and this tendency is believed to be more prominent in females, resulting in greater spatial cohesion among related females [15, 16]. The smaller recall scores of the females may be rooted in such behaviors, with smaller individual variations resulting in greater difficulties in IR.

Individual-specific patterns (i.e. personality) shown in this study have allowed researchers to further refine ecological and evolutionary models for Japanese macaques (reviewed in [17]). Previous studies on animal movement have focused on group-level behavior, in relation to environmental contexts (e.g. season and distribution of foraging resources), group size, and other individual traits (e.g. sex, age, and social rank). In such studies, individual variations in movement were typically treated as noise; however, recent developments in biologging technology and machine learning (represented by the BLE beacons and the neural network analysis adopted here) have enabled fine-grained recording and non-parametric analysis of individual behavior. Accordingly, there is a growing interest in more detailed models in movement ecology, representing both individual- and group-level movements of social animals as well as their interactions (reviewed in [18]). This study may help to develop such detailed models, providing empirical data to explore such complex relations.

## Acknowledgements

We appreciate the animal care by our technicians and research assistants as well as their suggestions and supports for this project. In particular, we wish to express special thanks to Norihiko Maeda, Mayumi Morimoto, and Takayoshi Natsume. We also acknowledge the profound comments and suggestions from Zin Arai and Hiroshi Takeuchi regarding the analysis. This study was performed under the Cooperative Research Program at KUPRI (2018-C-27, 2019-B-27).

## Author contributions

Project organization: IM HK; Animal arrangements: HK SA NSH; Apparatus building: AT IM HK; Data acquisition: AT IM HK; Animal cares: AK AT IM HK; Data management; HK AT TM; Computational modeling: TM; Manuscript writing: IM HK TM.

