## Supporting Information. for "Animals exhibit personality traits in their movement: a case study on location trajectories of primates"

### S1 Illustration of the Macaque Tracking System

Figure S1.1a illustrates our macaque tracking system based on the Bluetooth® Low Energy beacons. Figure S1.1b shows the custom-made collar worn by the macaques, fitted with Bluetooth® Low Energy beacons.

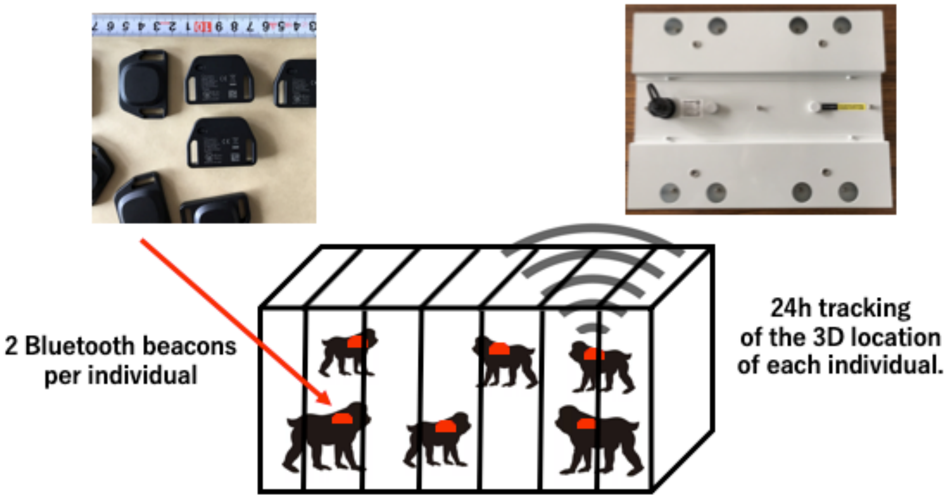


(a) System Illustration

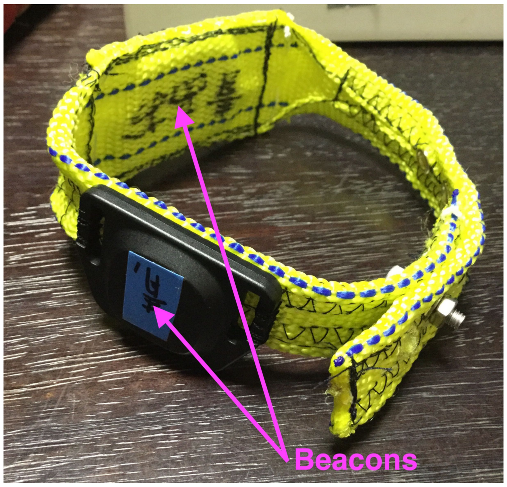


(b) Collar

Figure S1.1: (a) Illustration of the macaque tracking system. (b) The collar worn by the macaques. Two beacons were installed on each collar, indicated by purple arrows in the figure.

### S2 Neural Network

Our neural network received a 2-min trajectory of each individual's location ($= \mathbf{s}$, using samples from both beacons), and calculated a prediction of the individual corresponding to the input data, in the form of a probability distribution over the five individuals. Location data $(x_t, y_t, z_t)^{\mathrm{T}}$ were normalized so that each axis ranges between -1 and 1. Note that, while input data were defined by a chronologically fixed interval, their length varied because of an unstable sampling rate.[1] Accordingly, we adopted a recurrent neural network (RNN) to embed the time series data into a fixed-dimensional feature space. The particular implementation of the RNN was the bidirectional long short-term memory (LSTM, [1, 2]).

Figure S2.2 illustrates our network architecture. As described above, we first fed the time series input-denoted by the (normalized) coordinate vectors $(x_t, y_t, z_t)^{\mathrm{T}}$—to the bidirectional LSTM. We then concatenated the last hidden and cell states of both the directions and passed the concatenated vector to a multilayer perceptron (MLP). The MLP converted the LSTM-extracted features into unnormalized log odds of the individuals, and we finally obtained the normalized probabilities, $\mathbb{P}(I = 1 \mid \mathbf{s}), \ldots, \mathbb{P}(I = 5 \mid \mathbf{s})$, by taking the softmax function of the odds.

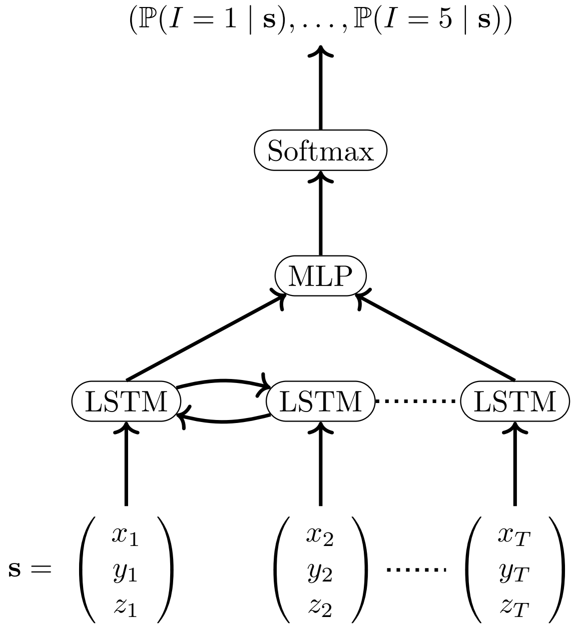


Figure S2.2: The architecture of the neural network predicting the individual's identity from their movement.

The free parameters of the network were set as follows: the bidirectional LSTM and the MLP had a single hidden layer; the number of hidden units of the LSTM and MLP were both 128; and the non-linearity of the MLP was tanh.

The network was trained by the stochastic gradient descent algorithm for 30 epochs with the cross-entropy loss. The learning rate was initially set as 1.0 and was multiplied by 0.1 after every epoch in which validation loss was not improved. The gradient norms were clipped at 1.0 to avoid the explosion. No dropout or momentum were introduced.

### S3 Data Size per Individual and Data Type

We randomly split the 175,494 time series into three types: 81% (142,149 series) were used for the training, 9% (15,795 series) for the validation, and 10% (17,550 series) for the test. Table S3.2 provides detailed information about the data size for each individual and data type.

### S4 Mathematical Definition of the Scores

The scores of the held-out test data were mathematically defined as follows:

[1] An unstable sampling rate could have put an extra burden on the network, requiring the network to predict the duration between two samples in the interval. Accordingly, we also trained the same network feeding the time information (normalizing 24 h time into a range on a scale of ?1 to 1) together with the location trajectory. However, this adjustment did not improve model predictions.

| | Data Type | | |
|---|---|---|---|
| Individual | Training | Validation | Test |
| Female 1 | 28394 | 3187 | 3528 |
| Female 2 | 28366 | 3129 | 3576 |
| Female 3 | 28521 | 3147 | 3444 |
| Male 1 | 28507 | 3089 | 3505 |
| Male 2 | 28361 | 3243 | 3497 |

Table S3.2: Data size (# of time series) per individual and data type.

| Target Individual | Score Type | Value | 95%CI | vs. Chance |
|---|---|---|---|---|
| Female 1 | Precision | 0.336160 | [0.318093, 0.354553] | $p < 0.001$ |
| | Recall | 0.247166 | [0.232932, 0.261500] | $p < 0.001$ |
| | F1 | 0.284874 | [0.270027, 0.299694] | $p < 0.001$ |
| Female 2 | Precision | 0.398670 | [0.380818, 0.416638] | $p < 0.001$ |
| | Recall | 0.318512 | [0.303320, 0.333883] | $p < 0.001$ |
| | F1 | 0.354112 | [0.339062, 0.369068] | $p < 0.001$ |
| Female 3 | Precision | 0.357045 | [0.339713, 0.374542] | $p < 0.001$ |
| | Recall | 0.301684 | [0.286374, 0.316973] | $p < 0.001$ |
| | F1 | 0.327038 | [0.312079, 0.341896] | $p < 0.001$ |
| Male 1 | Precision | 0.410964 | [0.397183, 0.424765] | $p < 0.001$ |
| | Recall | 0.577461 | [0.560996, 0.593891] | $p < 0.001$ |
| | F1 | 0.480190 | [0.466872, 0.493315] | $p < 0.001$ |
| Male 2 | Precision | 0.342402 | [0.328257, 0.356653] | $p < 0.001$ |
| | Recall | 0.417501 | [0.401249, 0.433796] | $p < 0.001$ |
| | F1 | 0.376240 | [0.362502, 0.389878] | $p < 0.001$ |

Table S5.3: The precision, recall, and F1 scores of the model predictions for each of the subject macaques. The statistical significance was evaluated against the chance level (= 0.2), from 100,000 bootstrapped samples.

- The accuracy score:
  $N^{-1} \sum_{j=1}^{N} \mathbb{1}[\operatorname{argmax}_i \mathbb{P}(I = i \mid \mathbf{s}^{(j)}) = i_{\mathrm{GT}}^{(j)}]$
- The F1 score of the detection of each individual $i$:
  - Precision: $\frac{\sum_{j=1}^{N} \mathbb{1}[i_{\mathrm{GT}}^{(j)}=i \wedge \operatorname{argmax}_{i'} \mathbb{P}(I=i' \mid \mathbf{s}^{(j)})=i]}{\sum_{j=1}^{N} \mathbb{1}[\operatorname{argmax}_{i'} \mathbb{P}(I=i' \mid \mathbf{s}^{(j)})=i]}$
  - Recall: $\frac{\sum_{j=1}^{N} \mathbb{1}[i_{\mathrm{GT}}^{(j)}=i \wedge \operatorname{argmax}_{i'} \mathbb{P}(I=i' \mid \mathbf{s}^{(j)})=i]}{\sum_{j=1}^{N} \mathbb{1}[i_{\mathrm{GT}}^{(j)}=i]}$
  - F1 score: the harmonic mean of the precision and the recall.

where $j$ indexes the test data, and $i_{\mathrm{GT}}^{(j)}$ is the ground truth individual responsible for the location trajectory $\mathbf{s}^{(j)}$ of the $j$-th test data.

### S5 Precision, Recall, and F1 scores

Table S5 reports the precision, recall, and F1 scores of the model predictions for each subject, supporting the results reported in Table 1.
